# A mathematical model for electrical activity in pig atrial tissue

**DOI:** 10.1101/2021.10.18.464761

**Authors:** Víctor Peris-Yagüe, Tony Rubio, Funsho E. Fakuade, Niels Voigt, Stefan Luther, Rupamanjari Majumder

## Abstract

Atrial fibrillation (AF) is the most common sustained form of cardiac arrhythmia occurring in humans. Its effective treatment requires a detailed understanding of the underlying mechanisms at the genetic, molecular, cellular, tissue and organ levels. To study the complex mechanisms underlying the development, maintenance and termination of cardiac arrhythmias, we need preclinical research models. These models range from *in vitro* cell cultures to *in vivo* small and large animal hearts. However, translational research requires that the results of these animal experiments are understood in the context of human subjects. Currently, this is achieved through simulations with state-of-the-art mathematical models for human and animal heart tissue. In the context of AF, a model that is extensively used by experimentalists, is that of the pig atria. However, until now, an ionically detailed mathematical model for pig atrial tissue has been lacking, and researchers have been forced to rely on mathematical models from other animal species to understand their experimental observations.

In this paper, we present the first ionically detailed mathematical model of porcine atrial electrophysiology. To build the model, we first fitted experimental patch-clamp data from literature to describe the individual currents flowing across the cell membrane. Later, we fine-tuned the model by fitting action potential duration restitution (APDR) curves for different repolarisation levels. The experimental data for the APDR studies was produced in N. Voigt’s lab.

We extended our model to the tissue level and demonstrated the ability to maintain stable spiral waves. In agreement with previous experimental results, our model shows that early repolarisation is primarily driven by a calcium-mediated chloride current, *I*_*ClCa*_, which is completely inactivated at high pacing frequencies. This is a condition found only in porcine atria. The model shows spatiotemporal chaos with reduced repolarisation.

**Author summary:** State-of-the-art mathematical models of cardiac electrophysiology play an important role in bridging the gap between animal research conducted in the laboratory and preclinical research being considered for translation into the clinic. Using computer simulations, these models enable detailed studies of the behaviour of ion channels and ion transfer at the cellular level, the propagation of electrical waves at the tissue level and the visualisation of the excitation pattern within the heart wall at the organ level. Thus, they contribute to a better understanding of the mechanisms underlying cardiac arrhythmias.

Here, we present the first ionically detailed mathematical model for porcine atrial electrophysiology. The individual membrane currents were modelled by fitting experimental data obtained from literature. The overall electrical response of the tissue was adjusted by fitting action potential duration restitution (APDR) curves obtained from in-house patch-clamp measurements. Our model accounts for an early repolarisation phase of the AP that is primarily Ca^2+^-dependent, a feature that is consistent with experiments and is identified to be unique to pigs. In extended media, our model is capable of sustaining stable spiral waves, and spatiotemporal chaos, when the repolarisation reserve is reduced.

## Introduction

Sudden cardiac death (SCD) is the most common cause of morbidity in industrialised countries [1–3]. Although most cases of SCD have been found to occur in patients with pre-existing coronary heart disease, a significant proportion of cases have been reported, which identify cardiac arrhythmias as the fatal precursor [4, 5]. Cardiac arhhythmias are electrical rhythm disorders that occur in the heart as a result of abnormal generation and/or propagation of electrical signals [8, 9]. The remediation of these disorders, requires the development of a detailed understanding of the origin, development and control over these abnormalities. There is general consensus that certain lethal types of cardiac arrhythmias, namely atrial fibrillation (AF), ventricular tachycardia (VT) and ventricular fibrillation (VF), manifest at the organ level as irregular, often reentrant, propagation of electrical waves through cardiac tissue [6]. Nonlinear dynamicists refer to these waves as spiral (in two dimensions) and scroll (in three dimensions) waves of electrical activity.

Despite the established concept that removing electrical spiral and scroll waves from the heart is essential for restoration of normal cardiac function, state-of-the-art technology to accomplish the same, remains suboptimal. This is because arrhythmias are very complex and multi-faceted diseases, which can have their roots at the organ-, tissue-, cellular-, molecular- or even genetic level. To understand how the origin affects the evolution of the disease or the specific mechanisms underlying arrhythmia control, it is important to conduct extensive research on diverse arrhythmia models. However, most arrhythmia research is conducted on small animal models, because of their larger availability, ease of maintainance in the laboratory environment and relative simplicity, compared to large animal, or human tissue models [7, 10]. Unfortunately, the problem with using small animals for experimentation is that the physiology and genetic makeup is inherently different from that found in large animal systems [10, 11]. This makes it difficult to transfer results obtained from small animal studies to the large animal models, leading to the creation of a broad gap between research that is ongoing in the laboratory and that which is approved for clinical translation. Mathematical models help to bridge this gap by transfering the knowledge obtained from large animal studies into ‘virtual human’ models, which can then be investigated at length without risking a patient’s life.

In order to transfer the results obtained from animal experiments as accurately as possible to research on humans, scientists use large animals whose heart anatomy and electrophysiology are similar to humans [11]. A particular example is the pig heart model [12–15]. It is used extensively for *in vivo* studies of cardiac arrhythmias because it resembles the human heart in size, anatomy and general electrophysiological properties. The pig atrial model is used to study atrial fibrillation (AF), the most common sustained form of arrhythmia that occurs in humans. However, it is a major challenge for scientists to compare the results of these studies with those in humans, because a detailed mathematical model describing the true electrophysiology of the pig, which can be used to bridge the aforementioned gap of knowledge transfer, is currently lacking.

In this study, we present the first detailed mathematical model of the pig atria, based on experimental patch-clamp data from literature and our own restitution experiments on pig atrial tissue. In two dimensions, we demonstrate its ability to produce and sustain stable meandering spiral waves. We characterise the spatiotemporal meander pattern, report the dominant frequencies and the effect of system size on the stability of the spiral pattern. In particular, we highlight some fundamental differences in the role of Cl^−^ currents and Ca^2+^ dynamics in the early repolarisation phase of AP in pigs with respect to humans, a crucial difference to be accounted for in the translatability of results, from pigs to humans, in future *in vitro* and *in vivo* experiments. We further go on to propose a model for AF using an altered set of parameters that allows us to have a state of electric turbulence in pig atrial tissue.

## Results

We developed a mathematical model for a native atrial cardiomyocyte, isolated from the excised atrium of a healthy adult pig. The equivalent electrical circuit representing the cell membrane is shown in Fig. 1.

**Fig 1.**
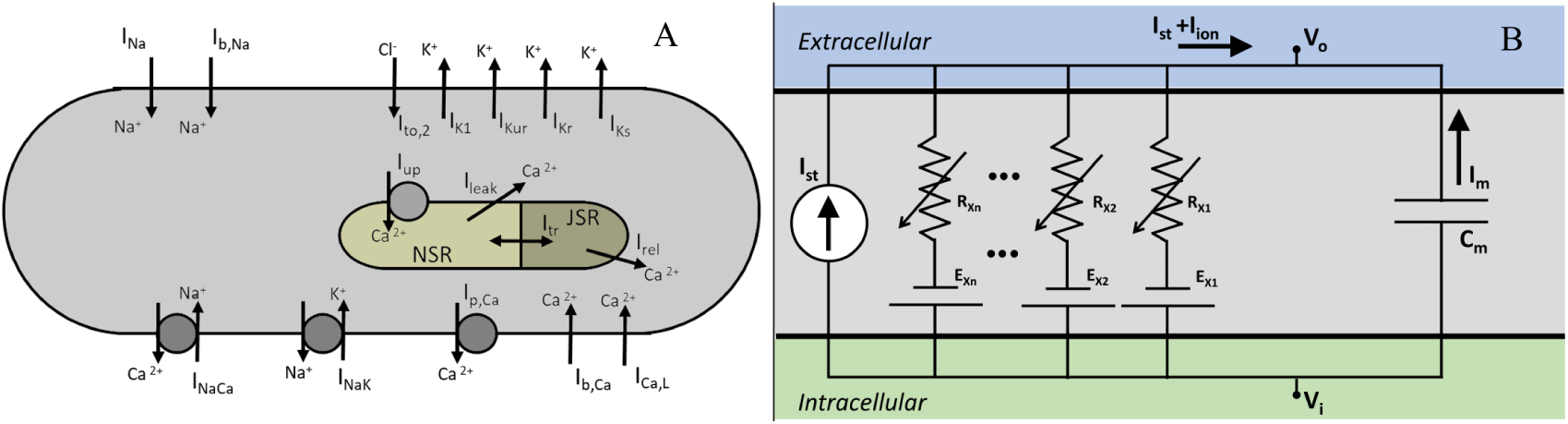
A. Schematics of a pig atrial cardiomyocyte model, showing transmembrane currents and the basic structure of the *Ca*^2+^ dynamics. B. Electrical circuit equivalent of the cell.

It consists of a membrane capacitance, *C*_*m*_, connected in parallel with several nonlinear conductances (*G*_*X*_) and batteries (*E*_*X*_). The net current (*I*_*ion*_) flowing across the cell membrane at any instant is the sum of individual currents flowing through the various branches of the circuit. Thus, the time-evolution of the transmembrane voltage *V* can be described using the following ordinary differential equation. (Eq. 1)

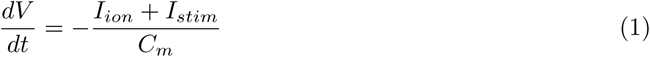

Here *I*_*stim*_ represents the external stimulus current that needs to be applied to the cell membrane to invoke an action potential (AP). We describe *I*_*ion*_ as a sum of 12 ionic currents (Eq. 2):

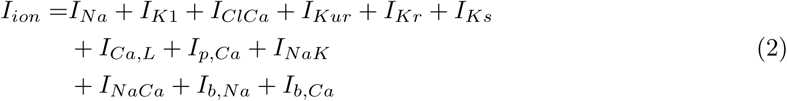

*V* is measured in millivolts (mV), time (*t*) in milliseconds (ms), *C*_*m*_ in picofarads (pF), and all currents in picoamperes per picofarad (pA/pF). All conductances *G*_*X*_ are measured in nanosiemens per picofarad (nS/pF), and intracellular and extracellular ionic concentrations ([*X*]_*i*_, [*X*]_*o*_) are expressed in milimolar (mM). The fast *Na*^+^ current is represented by *I*_*Na*_, the inward rectifier *K*^+^ current by *I*_*K*1_, ultrarapid rectifier *K*^+^ current is given by *I*_*Kur*_, rapid and slow delayed rectifier *K*^+^ currents by *I*_*Kr*_, *I*_*Ks*_, respectively, *L-Type Ca*^2+^ current by *I*_*Ca,L*_, *Ca*^2+^ pump current by *I*_*p,Ca*_, the Sodium-Potassium and Sodium-Calcium pump currents by *I*_*NaK*_, *I*_*NaCa*_, respectively, and the background *Na*^+^ and *Ca*^2+^ currents by *I*_*b,Na*_, *I*_*b,Ca*_, respectively. Uniquely in the pig atrial model, the transient outward current is represented only by a calcium-mediated chloride current, *I*_*ClCa*_.

The formulation of subcellular *Ca*^2+^ uptake and release by the sarcoplasmic reticulum (SR) is retained from the work of Luo and Rudy [18]. The three main currents involved are the *Ca*^2+^ uptake current, *I*_*up*_, the *Ca*^2+^ release current *I*_*rel*_ and the *Ca*^2+^ transfer current between the network SR (NSR) and junctional SR (JSR), *I*_*tr*_. The model also includes a leak current from the SR into the cytoplasm, *I*_*up,leak*_, as described by Courtemanche *et al*. in their model for the human atrial cardiomyocyte [19].

To invoke action potentials in tissue, we applied a stimulus current of 7*nA* for 4*ms*. The mathematical description of each ionic current is provided in the Appendix, together with a list of model parameters and initial values.

## Membrane currents

### Fast Sodium, *I*_*Na*_

We describe the fast *Na*^+^ current according to the Courtemanche-Ramirez-Nattel (CRN) model for human atrial cardiomyocytes [19], which uses a Hodgkin-Huxley type formulation (see Eq. 3):

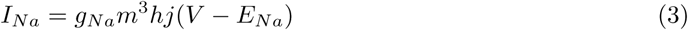

Here *m* is the activation gate; *h* and *j* are the two inactivation gates. In order to make the model pig-specific, we wanted to use Eq. 3 to fit experimentally obtained *I*_*Na*_ current-voltage (IV) characteristics and/or current traces from patch-clamp measurements. However, in the absence of these experimental data, we followed an alternative approach. Since *I*_*Na*_ is the only current that is active during the upstroke phase of an AP, it is considered to be responsible for the AP amplitude (APA) and maximum upstroke velocity 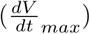 Thus, we used the complete model (considering all currents, pumps and exchangers) to fit experimentally obtained APA and 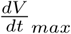 data, at various pacing frequencies, i.e., APA- and 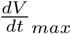 restitution.

Numerical fitting of these restitution data, using Eq. 3 to describe *I*_*Na*_ instructed us to apply the following adaptations: (*i*) raise the maximum channel conductance (*g*_*Na*_) by 80% with respect to humans; (*ii*) increase the time constant of activation (*τ*_*m*_) by a factor 1.7; and (*iii*) increase the time constant of inactivation (*τ*_*h*_, *τ*_*j*_) by a factor 2. The kinetics of the activation and inactivation gates are shown in Fig. 2A and B. With the applied modifications, the *I*_*Na*_ current traces turned out to be as in Fig. 2C and the IV curve, as shown in Fig. 2D.

**Fig 2.**
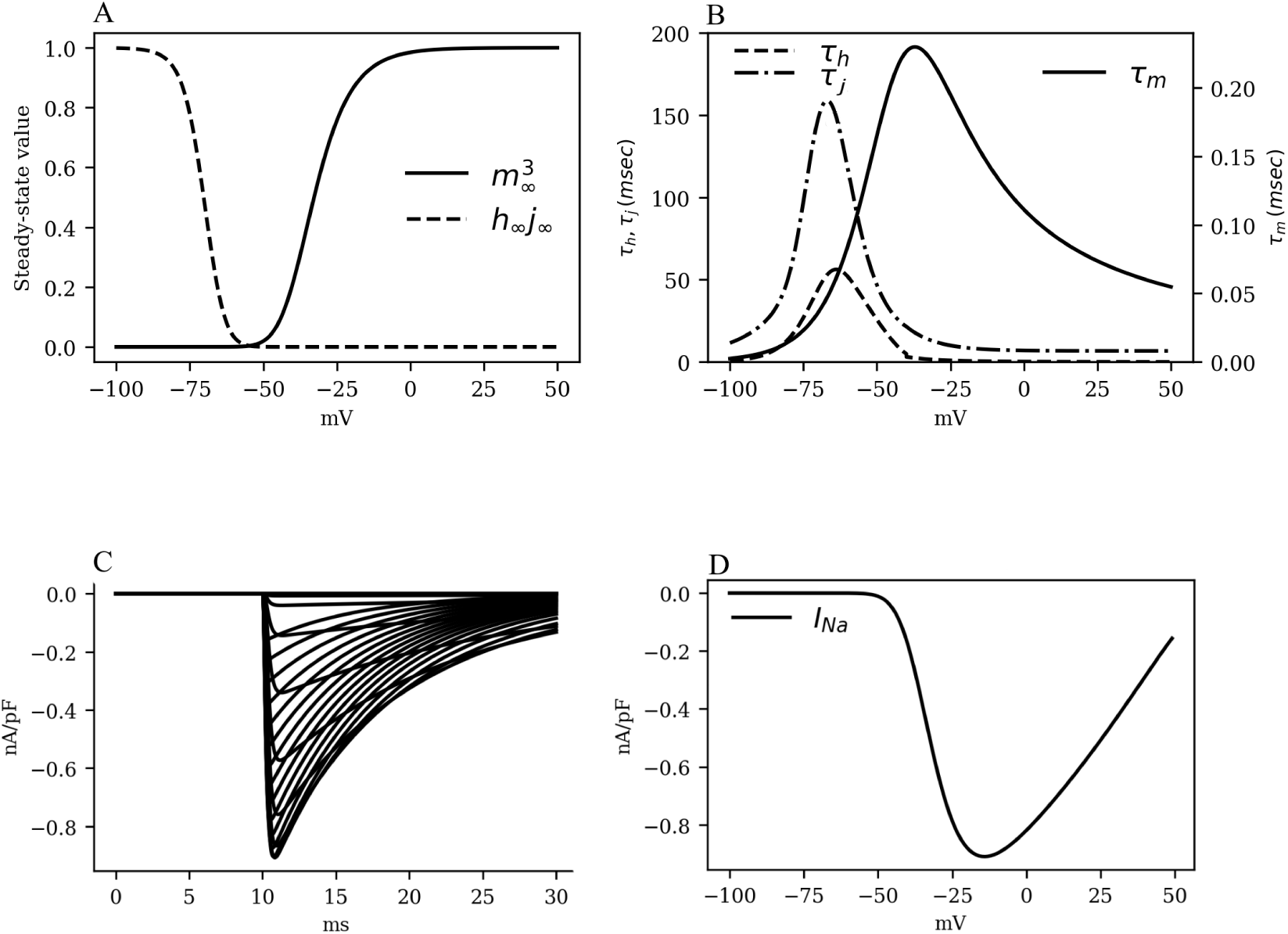
The kinetics of the fast *Na*^+^ current *I*_*Na*_. A. Activation and inactivation characteristics of the steady state gating variables *m* (raised to cubic power), *h* and *j* (combined as the product *h*_*∞*_*j*_*∞*_). B. Voltage dependence of the time constants for activation (*τ*_*m*_) and fast and slow inactivation (*τ*_*h*_ and *τ*_*j*_, respectively). C. Simulated traces of *I*_*Na*_. D. *I*_*Na*_ Current-Voltage characteristics.

### L-Type Calcium Current, *I*_*Ca,L*_

The L-type *Ca*^2+^ currrent (*I*_*Ca,L*_) was modelled according to previous literature [19, 21–23], based on the Hodgkin-Huxley formalism:

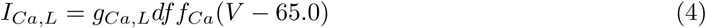

Here *g*_*Ca,L*_ is the channel conductance, *d* and *f* the voltage-gated activation and inactivation variables, respectively, and *f*_*Ca*_ is a calcium-mediated gating variable defined by:

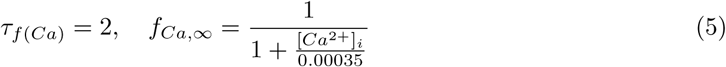

In particular, the gating behavior of *I*_*Ca,L*_ follows the human CRN model, with a +5 *mV* shift in the activation kinetics to decrease the activation window along with the overall *I*_*Ca,L*_ that is necessary for the fitting of restitution properties. The kinetics of L-type *Ca*^2+^ channel, as well as the IV characteristics of the *I*_*CaL*_ are shown in Fig. 3.

**Fig 3.**
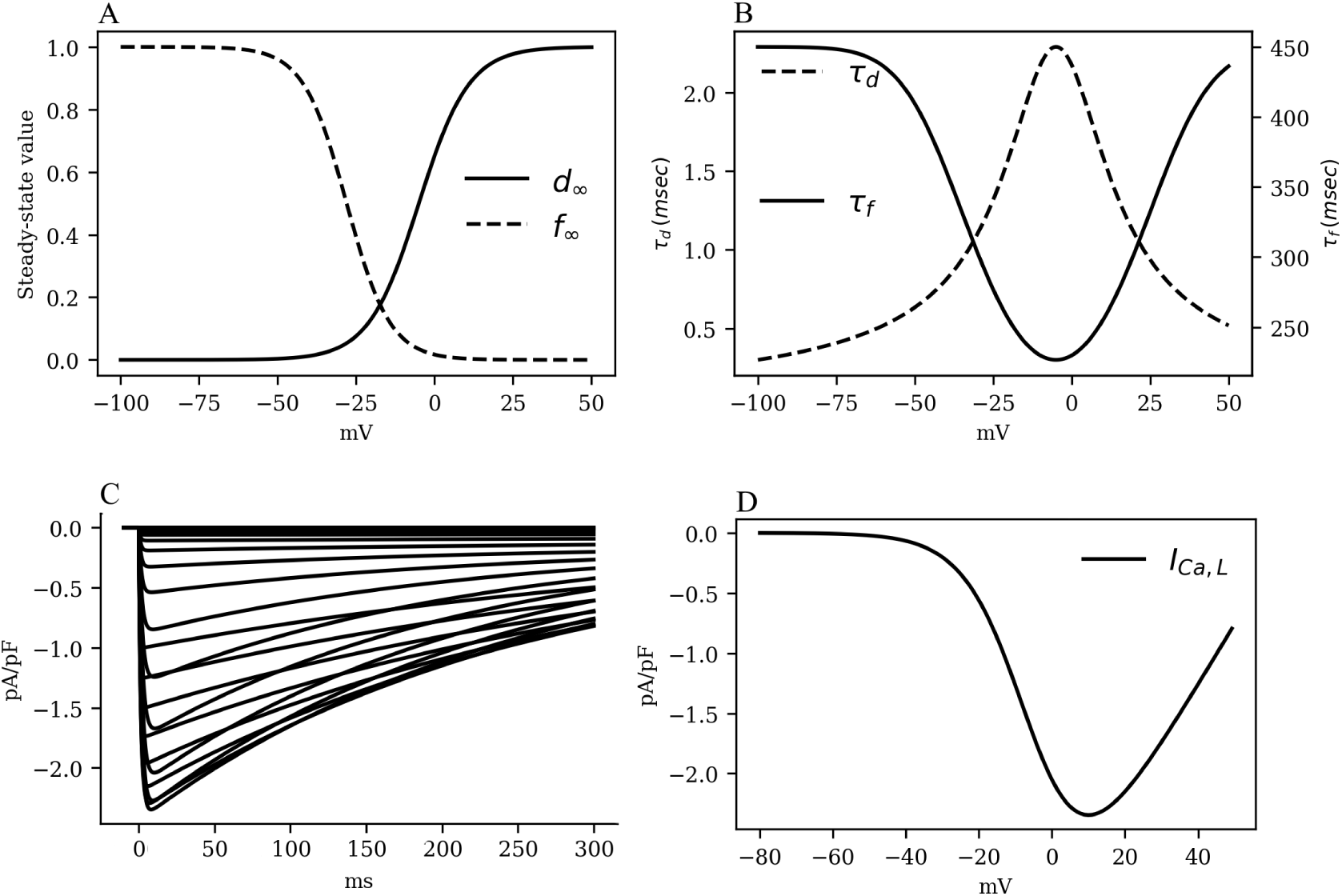
The kinetics of the fast L-type *Ca*^2+^ current *I*_*CaL*_. A. Activation and inactivation characteristics of the steady state gating variables *d* and *f*. B. Voltage dependence of the time constants for activation (*τ*_*d*_) and voltage-gated inactivation (*τ*_*f*_). C. Simulated traces of *I*_*CaL*_. Intracellular *Ca*^2+^ concentration was kept constant at [*Ca*^2+^]_*i*_ = 0.0001 *mM*. D. *I*_*CaL*_ Current-Voltage characteristics.

### Inward Rectifier Potassium Current, *I*_*K*1_

The *I*_*K*1_ is known to play a major role in determining the resting membrane potential (RMP) of excitable cardiac cells in many animal species. Given that our patch-clamp experiments on pig atrial tissue suggested a more depolarised (positive) RMP value than that reported in human atrial cardiomyocytes [19], we modified the parameters of the CRN *I*_*K*1_ formulation to make the current pig-specific.

To this end, (*i*) we shifted the reversal potential of *I*_*K*1_ by −5mV, as has been done previously in some large animal models, such as the sheep [20], (*ii*) we reduced the maximum channel conductance of *I*_*K*1_ by 9% relative to the human model [19], (*iii*) we modified the slope of activation of the *I*_*K*1_ IV curve by 10%; and (*iv*) We shifted the half-rise potential by +10 *mV*. Thus, *I*_*K*1_ in the pig atrial model is described according to Eq. 6. The IV curve for *I*_*K*1_ as shown in Fig. 4.

**Fig 4.**
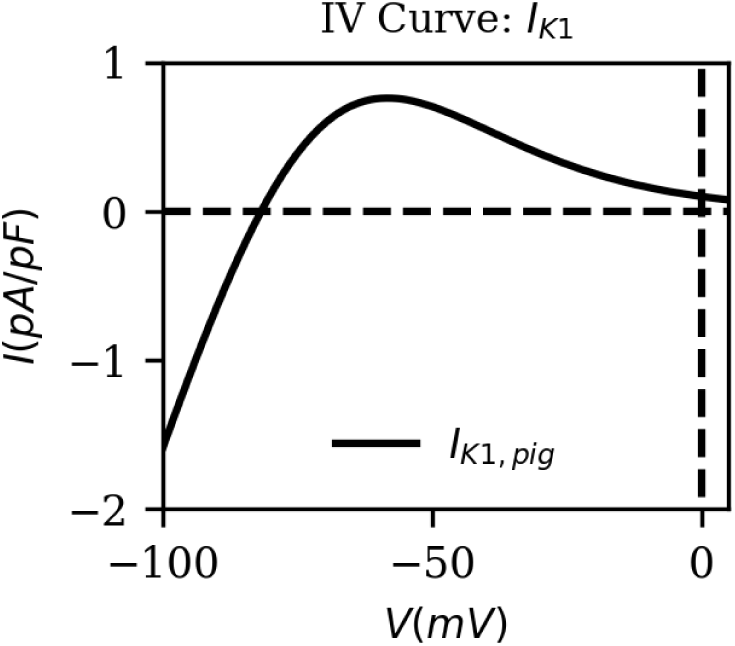
The current-voltage characteristic of the inward rectifier *K*^+^ current (*I*_*K*1_). The direction of flow of the current across the cell membrane reverses at *E*_*K*_ = − 81.76 *mV* which is close to our experimentally measured resting membrane potential.

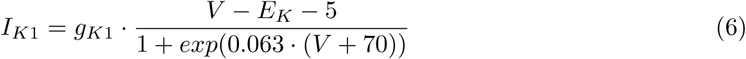

### Ultrarapid Potassium Current, *I*_*Kur*_

Recent work by Ehrlich *et al*. [25], indicates that pig atrial tissue exhibits a bi-exponential inactivation. Pandit *et al*. [24] developed a model for the ultrarapid *K*^+^ current, that reproduces the experimental data of Ehrlich *et al*. [25]. We used the formulation of Pandit *et al*. [24] to describe *I*_*Kur*_ in our model for the pig atrial tissue (see Eq. 7)

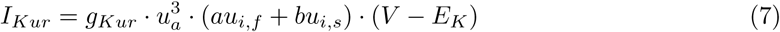

Here *g*_*Kur*_ is the channel conductance, *u*_*a*_ the activation gate, *E*_*K*_ the reversal potential of *K*^+^, and *u*_*i,f*_ and *u*_*i,s*_ the fast and slow inactivation components, respectively. (*a, b*) = (0.25, 0.75) are weights applied to the inactivation gates. It is interesting to note that the approach by Pandit *et al*. is similar to that of Aguilar *et al*. for the human atria. However, in the latter case, the inactivation of *I*_*Kur*_ is given by *u*_*i*_ = *u*_*i,f*_ *u*_*i,s*_ instead of a sum of the variables [26]. The conductance of *I*_*Kur*_ is described according to Eq. 8.

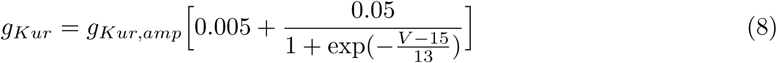

Here *g*_*Kur,amp*_ is an adjustable parameter whose value is determined during the final stages of model development (see sub-section Restitution Studies, for more details). Fig. 5A and B show the activation and inactivation kinetics of *I*_*Kur*_, whereas, a comparison between the current-voltage characteristics, as measured in experiments by Ehrlich *et al*. [25] and that produced using our model, is presented in Fig. 5C.

**Fig 5.**
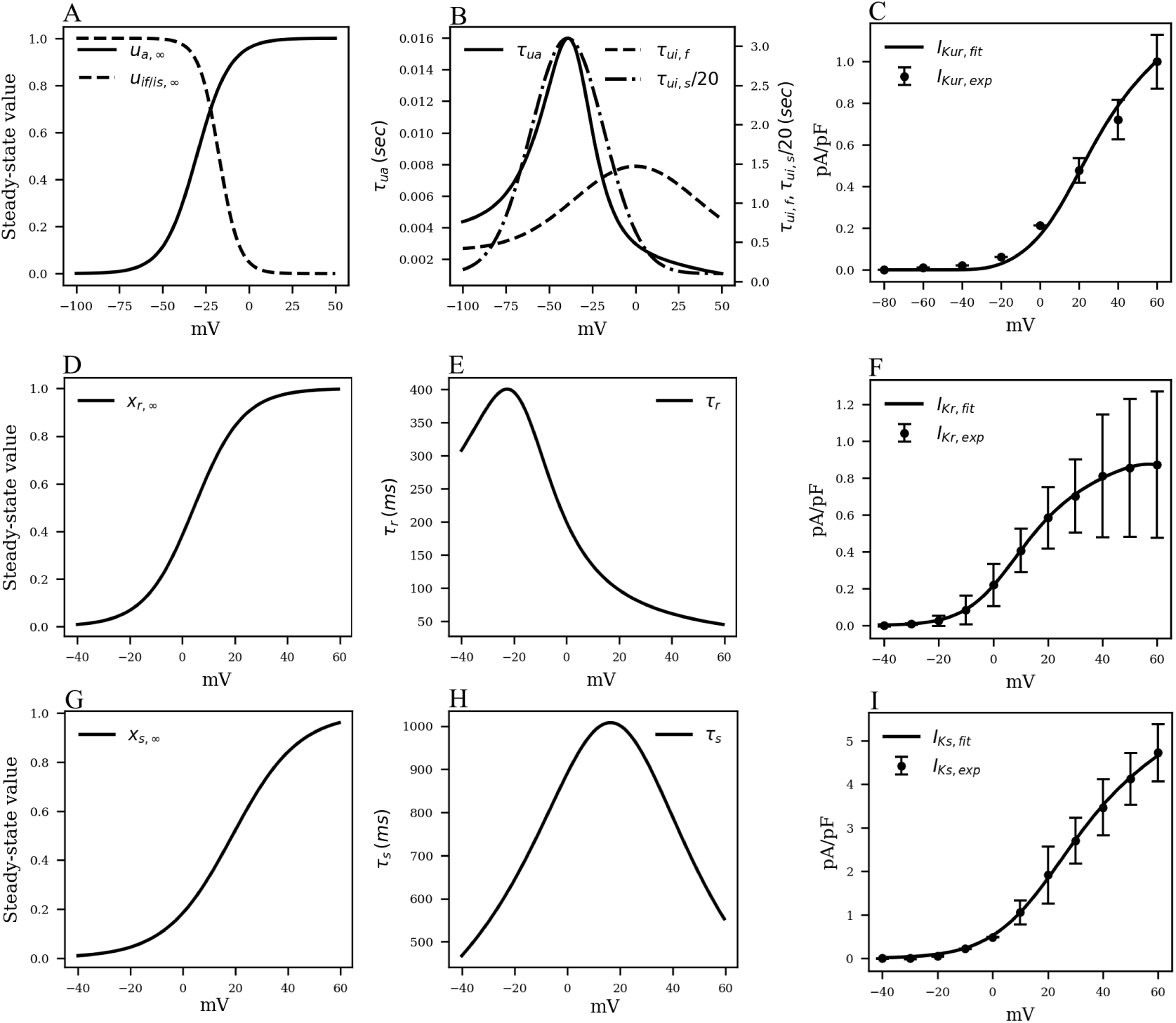
Channel kinetics and current-voltage characteristics of the rectifying *K*^+^ currents *I*_*Kur*_, *I*_*Kr*_ and *I*_*Ks*_. A. Voltage-dependence of steady-state activation (*u*_*s,∞*_) and inactivation (*u*_*if,∞*_ of *u*_*is,∞*_) probabilities of *I*_*Kur*_. B. Voltage-dependence of the time constants of activation (*τ*_*ui*_) and inactivation (*τ*_*uf*_ or *τ*_*us*_) of *I*_*Kur*_, reduced by a factor of 20. C. Comparison between the model-generated IV curve for *I*_*Kur*_ and the IV curve reported by Ehrlich *et al*. [25] based on experiments. D. Voltage-dependence of steady-state activation (*x*_*r,∞*_) probability of *I*_*Kr*_. E. Voltage-dependence of the time constant of activation (*τ*_*r*_) of *I*_*Kr*_. F. Comparison between the model-generated IV curve for *I*_*Kr*_ and the experimentally obtained IV curve from Li *et al*. [17].

### Rapid Delayed Rectifier Current, *I*_*Kr*_

The rapid delayed rectifier current (*I*_*Kr*_ was formulated similar to the original CRN model [19], but with altered half-rise voltage (*V*_1*/*2_), slope of the correction value, and the steady-state of the single gating variable, such that the obtained IV characteristic curve matches with the experimental IV curve of Li *et al*. [17]:

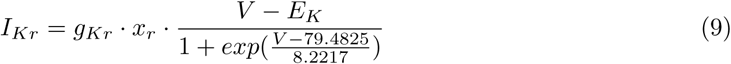

Initially, the conductance *g*_*Kr*_ was set at 0.0065 *pA/pF* to match the non-normalised IV curve. However, this value was tuned during the final stages of model development to match the pig atrial APD restitution curves at the tissue level. The steady-state value (*x*_*r,∞*_) of the gating variable *x*_*r*_ is described according to Eq. 10:

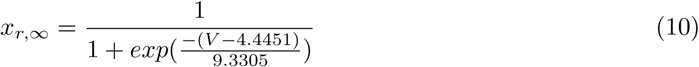

Note that, *V*_1*/*2_ of *x*_*r,∞*_ is shifted by +20*mV* relative to the CRN model, whereas, the slope of the *x*_*r*_ kinetic is slightly decreased [19]. The gating behavior of *x*_*r*_ is described in Fig. 5 D,E. Unavailability of sufficient experimental data led us to retain the temporal dynamics of the gating variable *x*_*r*_ from the CRN model [19]. Fig. 5 F shows the comparison between the experimental and simulated IV curves for *I*_*Kr*_.

### Slow Delayed Rectifier Current, *I*_*Ks*_

We retained the slow delayed rectifier current formulation from the original CRN model [19]:

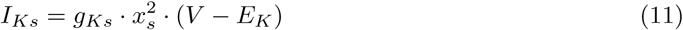

The maximum channel conductance *g*_*Ks*_ was adjusted to fit the restitution properties. The gating variable *x*_*s*_, and time constant *τ*_*s*_ are described according to Eqs. 12 and 13, respectively.

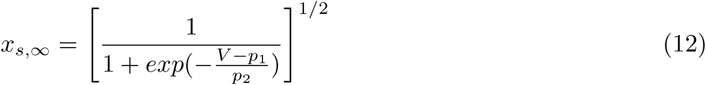

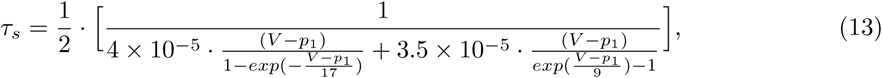

Here, parameters *p*_1_ and *p*_2_ have values 18.802 mV and 12.6475 mV, respectively, obtained by fitting experimental data from Li *et al*. [17]. The close resemblance of these values with those used in the human atrial tissue model [19] suggests that electrophysiologically, human atrial *I*_*Ks*_ and pig atrial *I*_*Ks*_ are very similar. Fig. 5 G and H show the steady state kinetic and time constant, respectively, of *I*_*Ks*_, whereas, the model-generated IV curve is compared with experimental data from Li *et al*. in Fig. 5 I.

### Transient Outward Current, *I*_*to*_

*I*_*to*_ in most species is composed of two components: a potassium current (*I*_*to*,1_) and a chloride current (*I*_*to*,2_, also referred to as *I*_*ClCa*_).

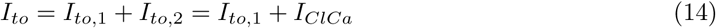

Although the presence of *I*_*ClCa*_ has been reported in multiple species and tissues [21, 27–31], including human atria [32, 33], it is generally observed that *I*_*to*,1_ forms the predominant component, scoring over *I*_*ClCa*_ in both strength and duration of activity, as a transient outward current [31, 32]. However, in a study by Li et al. [17], it was reported that pig atrial *I*_*to*_ is unique, in the sense that it is a completely calcium-driven chloride current [17, 34, 35]. Thus, in our model, we incorpotrated this feature by modelling *I*_*ClCa*_ according to Eq. 15, and experimental data from Li et al. [17].

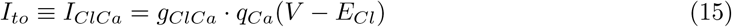

For the choice of formulation of *I*_*to*_ we considered various candidates [21, 29–31]. In the end, we decided to use Eq. 15, which is a formulation for *I*_*ClCa*_ in a canine atrial model [21]. The reason for choosing this formulation was that it allowed a fairly accurate reproduction of the bell shape of the IV curve and the same general upward trend present in the experimental data of Li *et al*. [17].

Here *g*_*ClCa*_ is the channel conductance, *E*_*Cl*_ the *Cl*^*−*^ reversal potential and *q*_*Ca*_ the sole gating variable of the channel, which follows the typical gating behavior of a Hodgkin-Huxley-type gating variable:

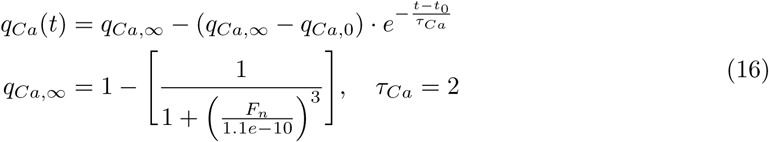

*F*_*n*_ is the flux of *Ca*^2+^ into the myoplasm. *F*_*n*_ shows a strong correlation with the sharp release of *Ca*^2+^ from the SR in the initial stages of AP (through the SR release current, *I*_*rel*_), giving *I*_*ClCa*_ the fast dynamics of a transient outward current. Also, the inactivation of *I*_*rel*_ gives *I*_*ClCa*_ a significant bell-shape in its IV curve, something universally observed in *I*_*ClCa*_ [21, 29, 36, 41]. In the case of our model, we shifted the inactivation of *I*_*rel*_ by +40 *mV* and increased the slope of inactivation to fit experimental results from Li *et al*. [17].

Fig. 6 shows the IV curve for *I*_*ClCa*_, as obtained using our model of the pig atrial tissue, overlaid on experimental data from Li et al. [17]. The simulated current activated earlier than in experiments, but with a good overall qualitative behavior.

**Fig 6.**
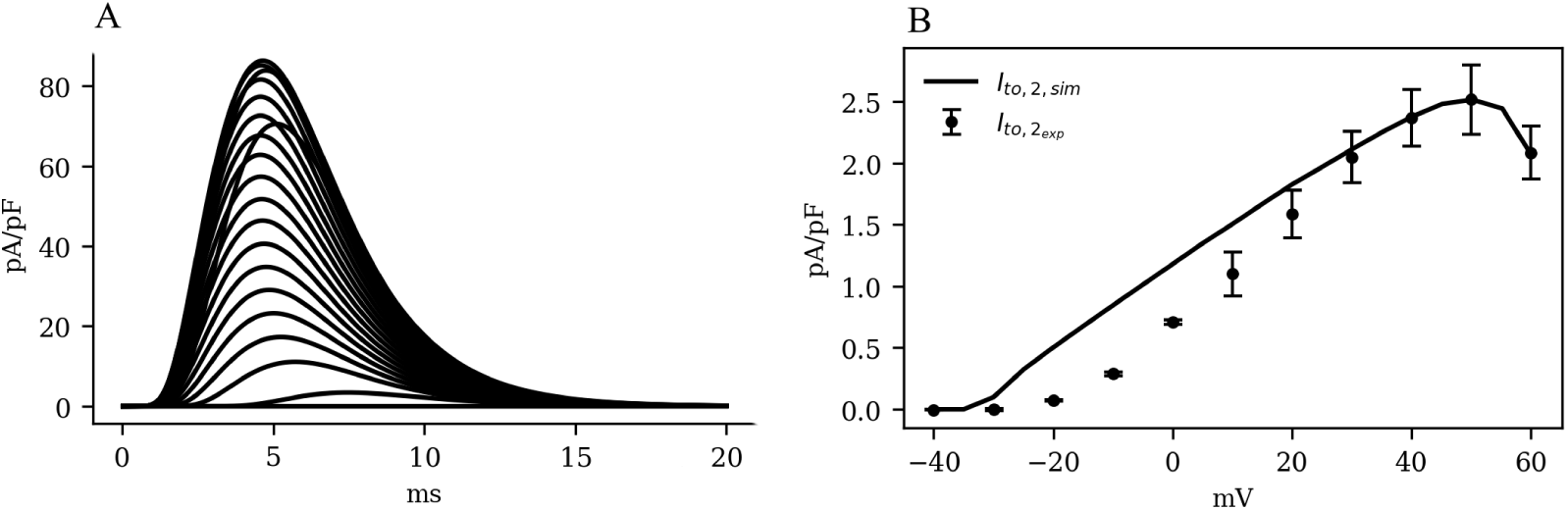
A. Simulated traces and (B) Current-voltage characteristics of *I*_*ClCa*_, showing experimental data (dots with error bars) from Li *et al*. [17], overlaid on the model-generated curve (solid). Note the significant bell-shape of the curve at high positive voltage values.

## Restitution Studies

### Action Potential Duration (APD)

The amount of time, during an action potential, when the membrane voltage of an excited cardiac cell is more positive than a chosen threshold, is called the action potential duration (APD) at that threshold. Typically, this threshold value is measured on the basis of degree of repolarisation of the cell membrane. Thus, APD_*X*_ refers to the amount of time during an AP, when the cell membrane is more than X% repolarised, or, less than X% depolarised.

When cardiac tissue is electrically stimulated using a train of pulses at a particular frequency, the morphology of the AP adapts to the applied pacing frequency. This reflects in the APA, RMP,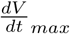 and APD values at all possible levels of repolarisation. Such studies are conducted to investigate the restitution behaviour of the model or the tissue sample. We performed patch-clamp experiments on pig atrial tissue to obtain APD restitution (APDR) data. We used these data to make final adjustments to the model, to perfect its electrical response to high frequency stimulation. In both experiments and simulations, pig atrial tissue was stimulated at 0.25, 0.5, 1, 2, 3, 4 *Hz*, and action potentials were recorded.

We adjusted model parameters to find the most optimal parameter set that simultaneously fit each of these restitution curves with minimal deviation from measurement. Specifically, we adjusted the maximum conductance values of several currents. The final selection of conductance values is listed in table 1.

**Table 1.** Conductance values and maximal currents after fitting restitution data.

| Conductance | Value (nS/pF) |
| --- | --- |
| $g_{K1}$ | 0.08218 |
| $g_{Na}$ | 13.9900 |
| $g_{Kur,amp}$ | 0.45539 |
| $g_{ClCa}$ | 0.15731 |
| $g_{Kr}$ | 0.01730 |
| $g_{Ks}$ | 0.0594 |
| $g_{Ca,L}$ | 0.06574 |
| Maximal Current | Value (pA/pF) |
| $I_{NaK,max}$ | 0.94935 |
| $I_{NaCa,max}$ | 2304 |

The overlays of our experimental and simulated data for each of the following parameters: APA, RMP,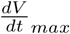, and APD_*X*_, for X=10, 20, 30, 40, 50, 60, 70, 80 and 90% repolaization are shown in Fig. 7.

**Fig 7.**
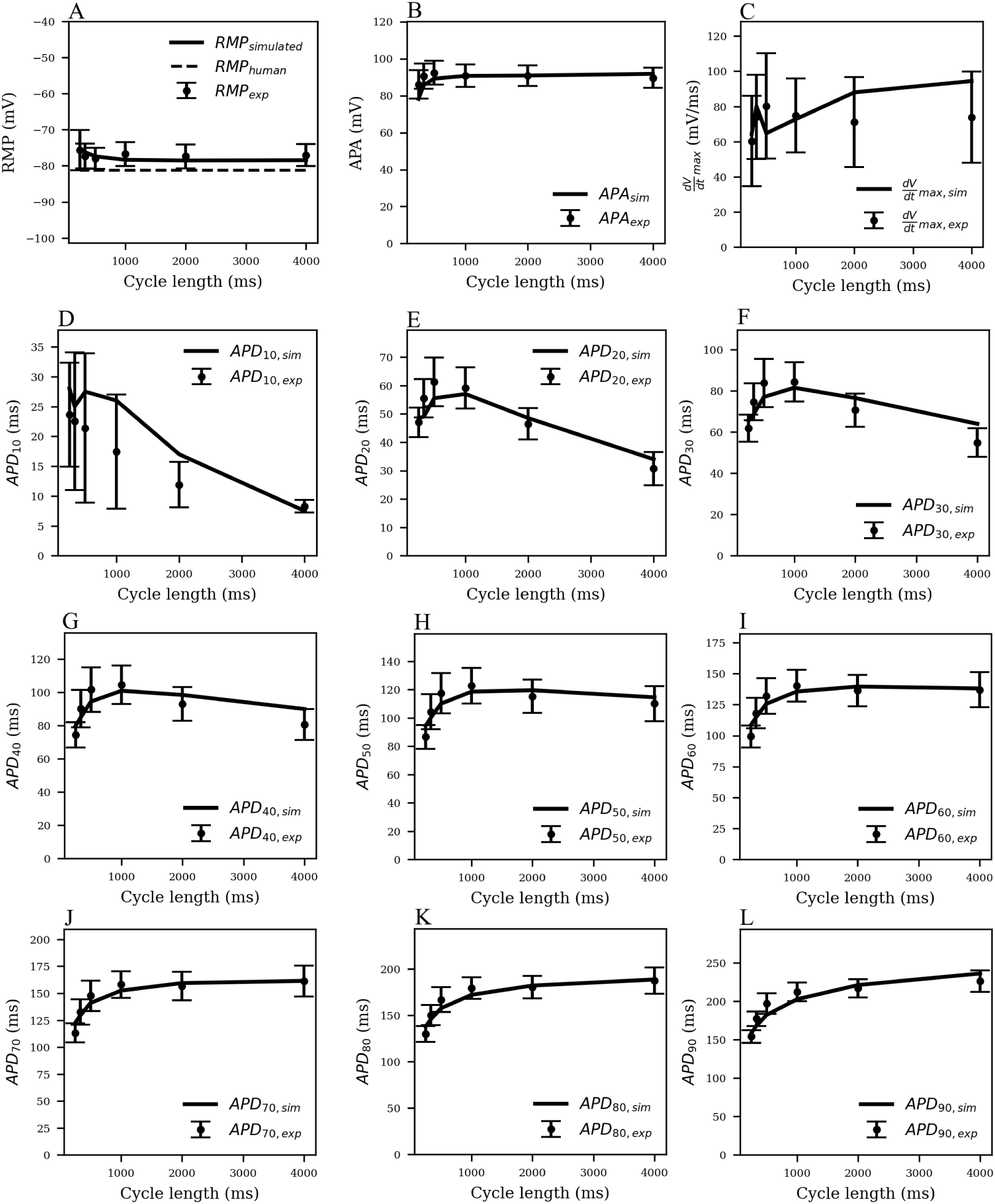
Restitution curves for (A) resting membrane potential (RMP), (B) action potential amplitude (APA), (C) maximum upstroke velocity 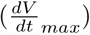, and (D-L) action potential duration (*APD*_*X*_), at *X* = 10, 20, 30, 40, 50, 60, 70, 80 and 90% repolarisation of the membrane voltage. Solid black lines indicate the model-generated data, whereas the markers (with error bars) represent our data obtained from patch-clamp experiments.

Note that pig APDR curves show an interesting feature that distinguishes the model from most other mammalian species that we know of. In the early stages of repolarisation (i.e., up to APD_50_), an overall downward trend is observed for stimulation cycle lengths greater than 1000 *ms* (see Fig. 7, D-H). This could be due to inactivation of *I*_*ClCa*_ at high pacing frequencies: slower and lesser calcium uptake causes a decrease in *I*_*ClCa*_, which significantly slows down initial AP repolarisation, causing an overall higher early APD at high pacing frequencies with respect to low-frequency pacing.

To test this hypothesis, we measured intracellular calcium flux and *I*_*ClCa*_ in simulated restitution experiments. Fig. 8 shows the resulting restitution curves for peak calcium flux and peak *I*_*ClCa*_ at different stimulation cycle lengths in the simulated model. A clear dependence of peak values on cycle length can be observed, and both quantities decrease significantly with a decrease in cycle length. This is indeed indicative of an initial AP repolarisation phase that is heavily dependent on calcium dynamics. To the best of our knowledge, this behavior is exclusive to the pig, and might have profound implications in the translatability of studies on arrhythmia control and termination from pigs to other species.

**Fig 8.**
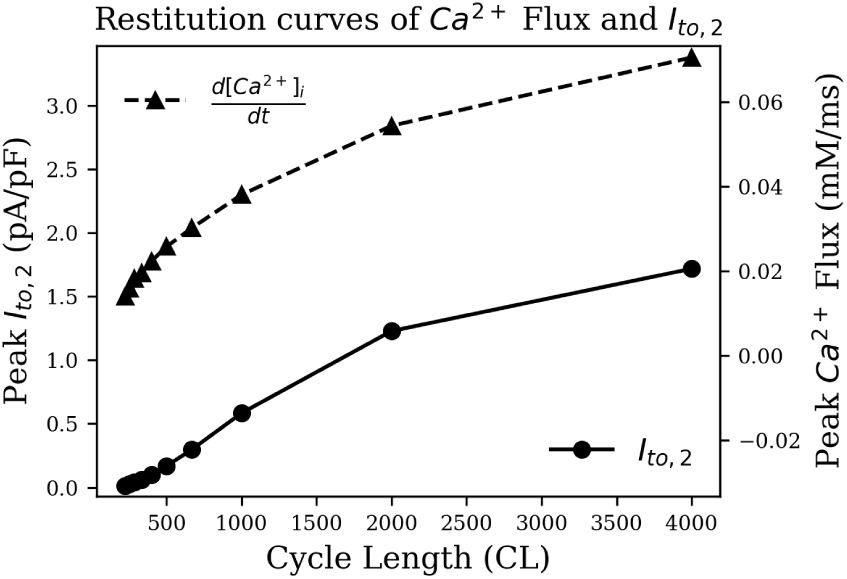
Restitution curve of peak intracellular calcium flux (dashed) and peak *I*_*ClCa*_ (solid).

To summarise, the pig atrial model is capable of reproducing experimentally recorded porcine APs at different pacing frequencies within experimental deviations (Fig. 9 A). The discrepancies between the two experimental traces shows the variable nature of electrophysiology, in particular in the atria [37], and the model is good at finding a compromise and reproducing an AP with average traits within experimental tolerances. Fig. 9 B, C and D show the temporal evolution of each of the transmembrane currents considered in Eq. 2. With this basic model, we now begin our study of electrical wave propagation at the tissue level.

**Fig 9.**
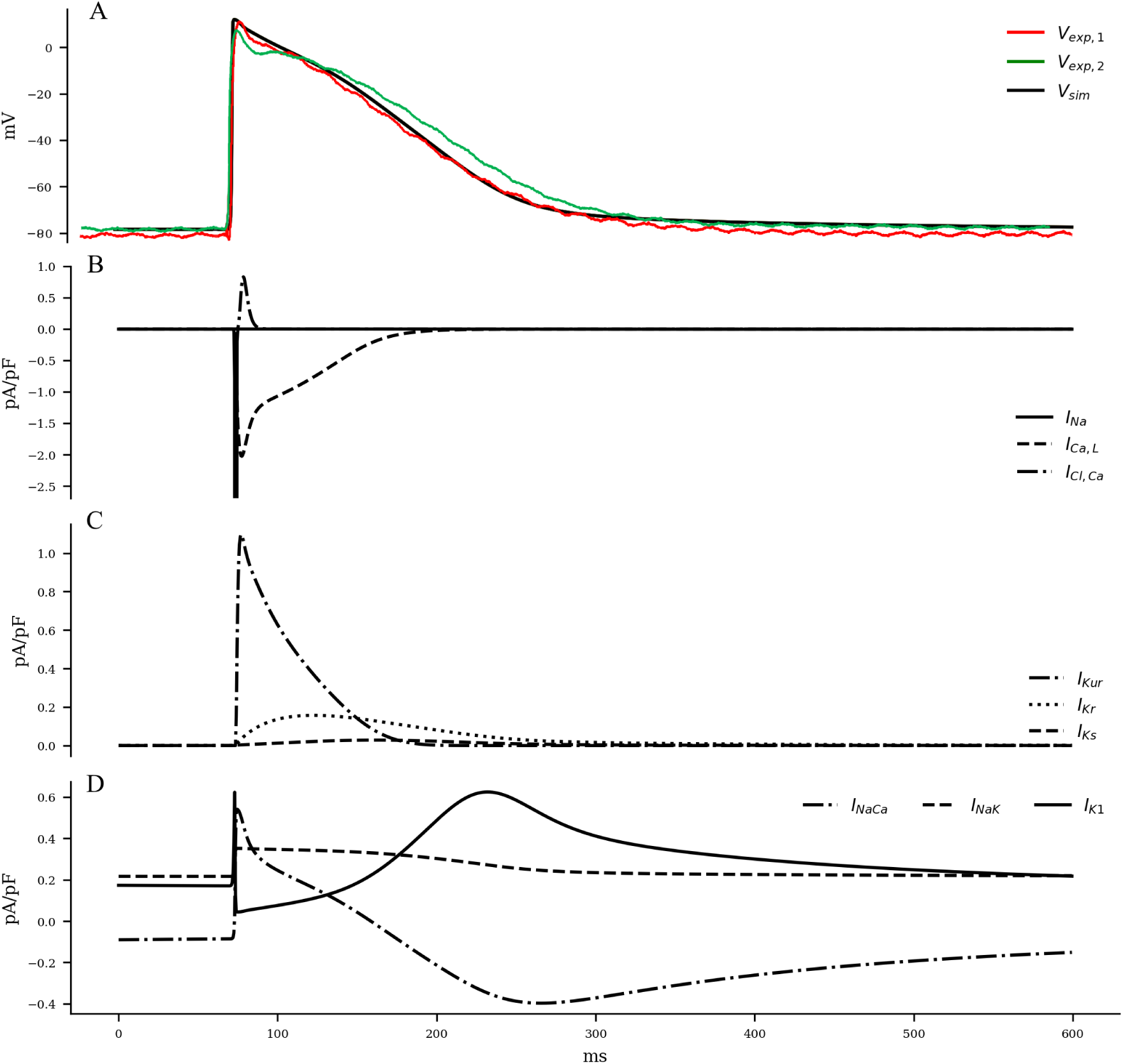
A. Voltage and (B-D) Current traces of a pig atrial action potential. The solid black line in shows results from our model simulation; green and red lines represent AP recordings from experiment. All Experiments and simulations were performed in tissue, but the voltage and current was measured at the single-cell level.

### Wave Propagation in 2D

Electrical stimulation of a quiescent 2D domain containing pig atrial cardiomyocytes leads to propagation of an excitation wave. Our studies confirm that at frequencies below 5 Hz, the paced waves propagate with uniform and identical wavefront and waveback conduction velocity. Electrical pacing at higher frequencies does not lead to 1:1 capture. This is because the effective refractory period of the cells is approximately 215 ms. Fig. 10 A shows snapshots of plane wave propagation through a 2D domain containing identical pig atrial cardiomyocytes. The simulated wavelength (WL, estimated as *WL* = (*t* ∣_*back*−40*mV*_ − *t* ∣_*wave front*_) × *CV*) and CV restitution curves are presented in Fig.10 B, C.

**Fig 10.**
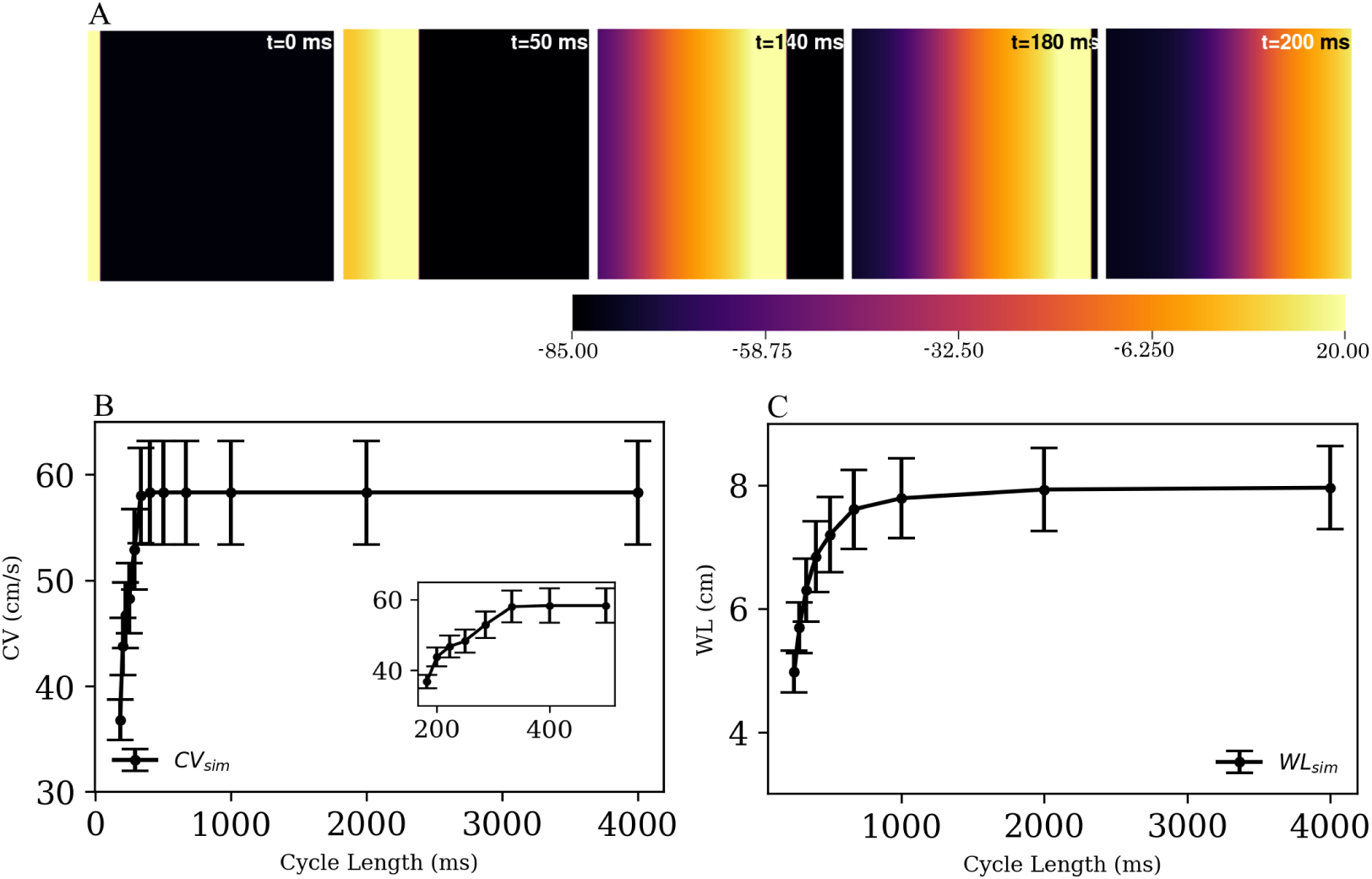
Propagation of a plane wave through simulated 2D pig atrial tissue. Panel A shows snapshots of the propagating wave at different times. (B) and (C) show the conduction velocity (CV) and wavelength (WL) restitution curves, as obtained from simulations.

## Spiral Waves in the Pig Atria

### Spiral Initiation

Using the reported parameter set in our 2D model, we produced a spiral wave that survived for more than 40 s of simulation time. To initiate a spiral wave in a domain containing 512× 512 grid points, we used the S1-S2 cross field protocol. We applied a line stimulus along the left edge of the domain to initiate a plane wave (S1) propagating towards the right (Fig. 10 A). As the waveback of the S1 wave crossed *x* = 256, a second stimulus (S2) is applied in the region *y* ≤ 256 (Fig. 11 A, *t* = 240 ms). This leads to propagation of the S2 wave in the region that has recovered from excitation. With time, as the wave S1 wave moves out of the domain, more excitable tissue becomes available and a spiral prototype is formed (Fig. 11 A, *t* = 280 ms). Fig. 11 panel A shows the spatiotemporal formation and evolution of the spiral.

**Fig 11.**
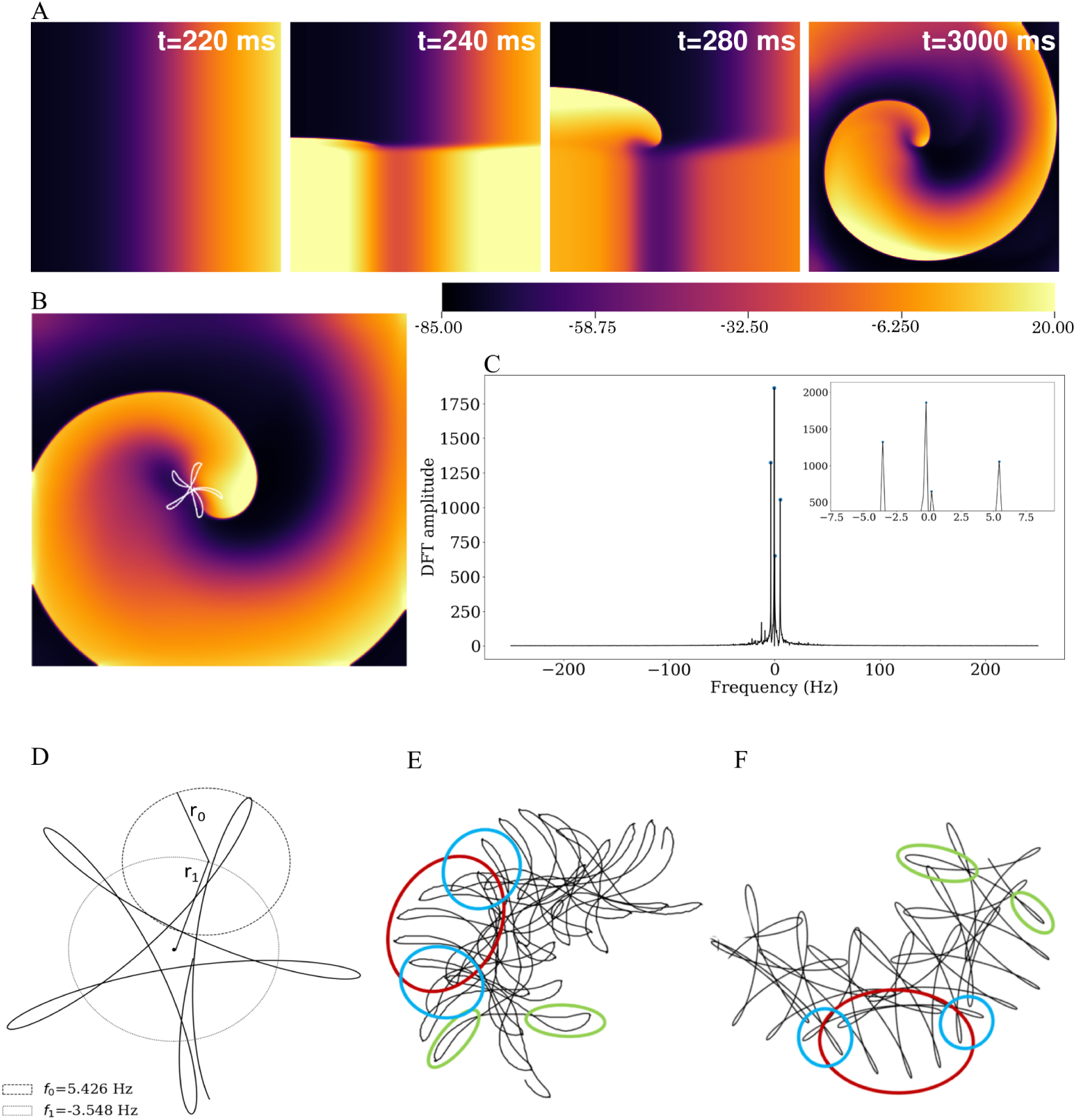
A. Formation and (B-F) characterisation of spiral waves in 2D pig atrial tissue. Here *t* = 0 is considered as the time instant at which the S1 wave is initiated (as in Fig. 10 A). B. Hypocycloid pattern of the spiral tip trajectory. C. Amplitude of the Discrete Fourier Transform (DFT) of the tip trajectory. The sampling frequency is *F*_*s*_ = 500 *Hz*, and the corresponding resolution Δ*f* = 0.2089 *Hz*. The inset shows the 4 main peaks of the DFT. D. Spiral tip trajectory reconstructed using the two significant peaks far from *f* = 0*Hz* from the Fourier Analysis at *f*_0_ = 5.426 *Hz* and *f*_1_ = −3.548 *Hz*. The same pattern is observed as in B. E. Tip trajectory recorded in a simulation with clockwise meander. The main features are: (*i*) thicker petals on the inside of the trajectory than on the outside (green), (*ii*) an overall counterclockwise movement of the pattern within (red) with periodic overlap of successive outer petals (blue). F. Reconstructed trajectory using the three dominant frequencies of the DFT (*f*_*meander*_, *f*_0_, *f*_1_). All three features discussed in E are qualitatively reproduced.

### Spiral Characterisation

The spiral wave in the pig atrial tissue model meanders with a shifting hypocycloidal trajectory. An analysis of the tip trajectory shows that the basic pattern contains 5 outward petals enclosing a center (see Fig. 11 B), which shifts in space at the end of every 5 rotational cycles. A Fourier analysis of the tip trajectory reveals the existence of 4 fundamental frequencies (see Fig. 11 C), of which *f*_0_ = 5.426 *Hz* and *f*_1_ = −3.548 *Hz* are the dominant ones. These contribute to the construction of the basic hypocycloidal pattern, through superposition of the two counter-rotating circular orbits at the given frequencies (the radii of the orbits are proportional to the heights of the peaks obtained from the Fourier Transform of the trajectory) (see Fig.11 D). The first sub-dominant frequency is responsible for the shifting of the center of the basic pattern of the tip trajectory, whereas the second sub-dominant frequency accounts for minor corrections to the global pattern. Although the resolution of DFT did not allow us to obtain exact values of these sub-dominant frequencies, an overall approximation with the use of a single sub-dominant frequency ≃ − 0.1 *Hz*, can result in the accurate reproduction of the most distinguishable features of the spiral tip movement (see Fig. 11 E, F).

### Alternans

A visual impression of the spatio-temporal distribution of membrane tension during spiral wave evolution (S1 Video) indicated the occurrence of wavelength fluctuations, as opposed to a constant, uniform wavelength observed during plane wave propagation.

Fig. 12 shows the restitution curves of APD_90_ the simulated spirals, both with respect to Cycle Length (CL) (Fig. 12 A) and Diastolic Interval (DI) (Fig. 12 B). Fig. 12 A shows the presence of alternans for cycle lengths in the range ≡ 165 − 215 *ms*. This is consistent with the restitution curve in Fig. 12 B, which focuses on the region with slope ≃ 1, a known predictor of the presence of alternans [39]. In a previous work Fakuade *et al*. [43] demonstrated the the occurrence of alternans at low stimulation frequencies in patients suffering from postoperative AF. Thus, our model can be used to develop useful insights into the origin and control of this alternans in pig atria.

**Fig 12.**
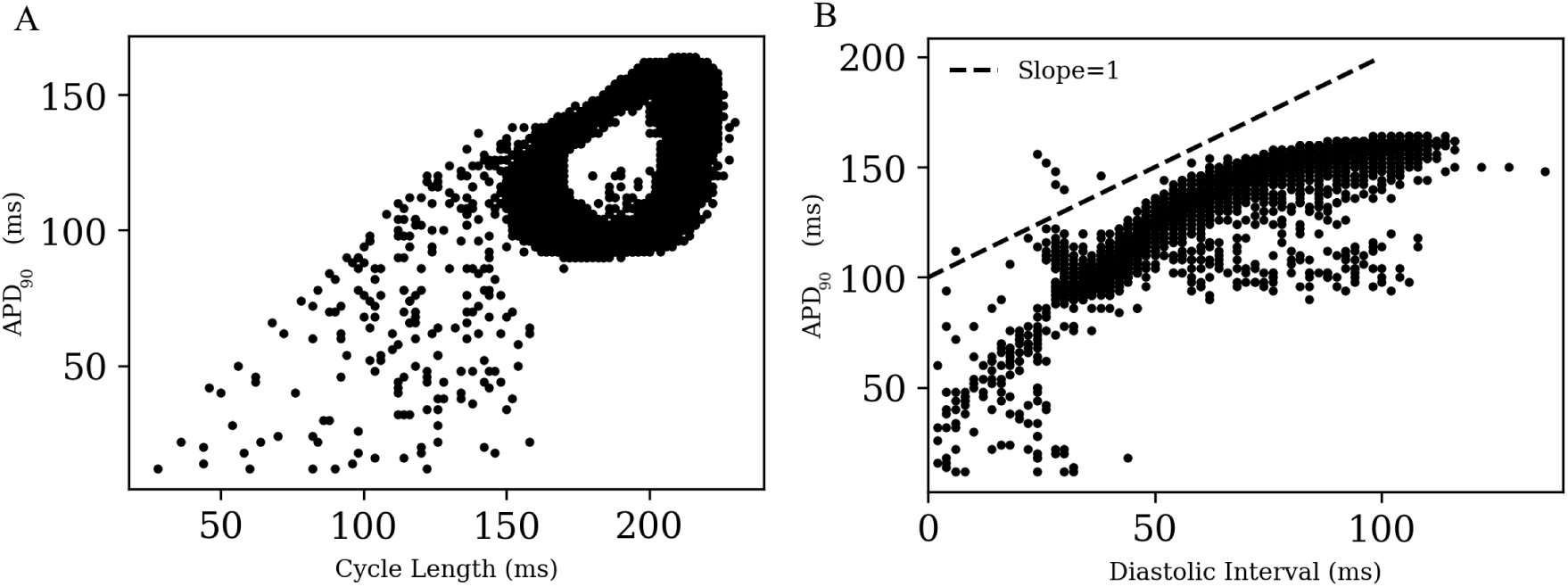
Restitution curves for *APD*_90_ in (A) a simulated spiral with respect to Cycle Length and (B) Diastolic Interval. The dashed line in (B) indicates the region of slope = 1

**Fig 13.**
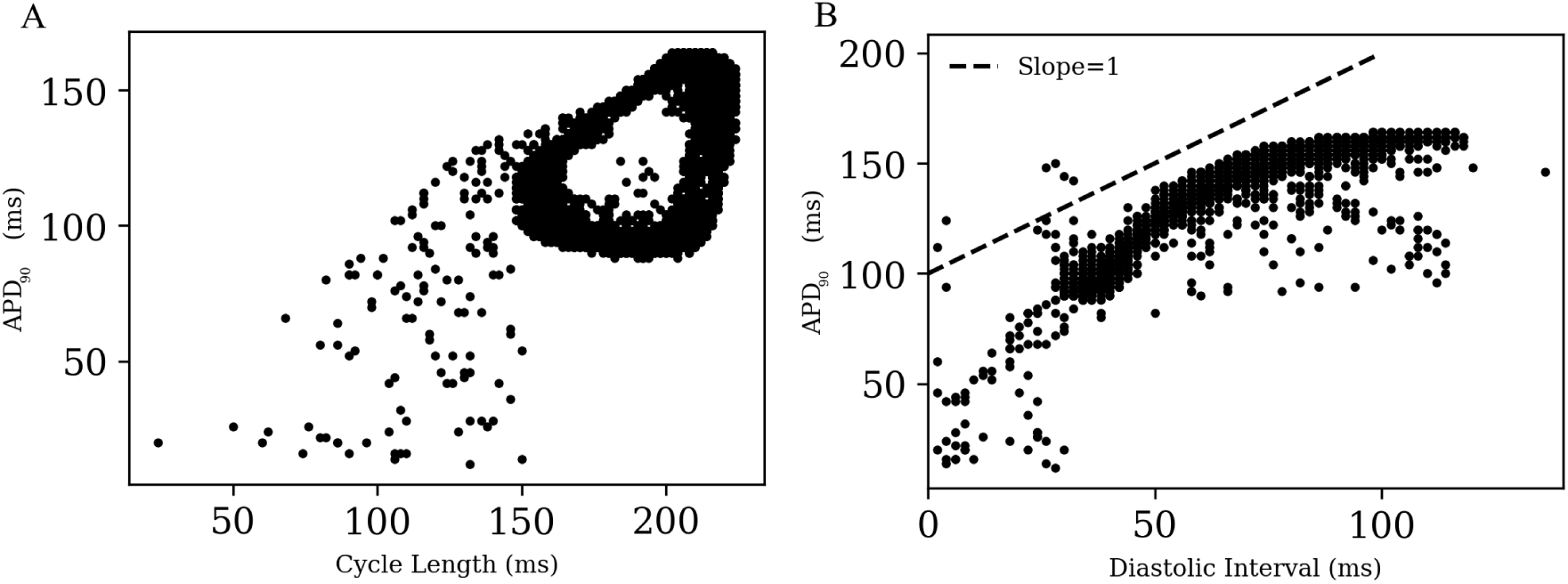
Restitution curves for *APD*_90_ in (A) a simulated spiral with respect to Cycle Length and Diastolic Interval, for a 90% reduction in *I*_*ClCa*_ with respect to control conditions. The dashed line in (B) indicates the region of slope = 1

To test if the unique current *I*_*ClCa*_ is responsible for alternans in the pig atrial model, we followed an approach that was first proposed by Gomis-Tena and Saiz [42]. Accordingly, we inhibited the *I*_*ClCa*_ (by 50% and 90% in two separate cases) in pig atrial model and re-initiated the spiral. However, unlike Gomis-Tena and Saiz [42], alternans continued to exist in our model. APs in the simulated spiral have a duration of at most ≃ 225 *ms*. Referring back to Fig. 8, we can see that *I*_*ClCa*_ is naturally already shut off at such small cycle lengths, and any further inactivation will obviously have a negligible effect on the behavior of the resulting spiral.

### Spiral Wave Breakup

Finally, we arrive at the most challenging question. Is it possible to use this model to study atrial fibrillation, with the spiral waves actually breaking up? The answer is, yes. The model does exhibit a state of sustained chaotic electrical activity in an altered parameter regime. Spiral wave breakup could be initiated by suppressing the repolarisation reserve. In particular, reducing the manimum channel conductance of *I*_*Kr*_ to 0.25x its original value could lead to a state characterised by more than six spiral waves. The spatiotemporal evolution of the spiral breakup state is demonstrated in Fig. 14 and in S2 Video of the Supporting Information.

**Fig 14.**
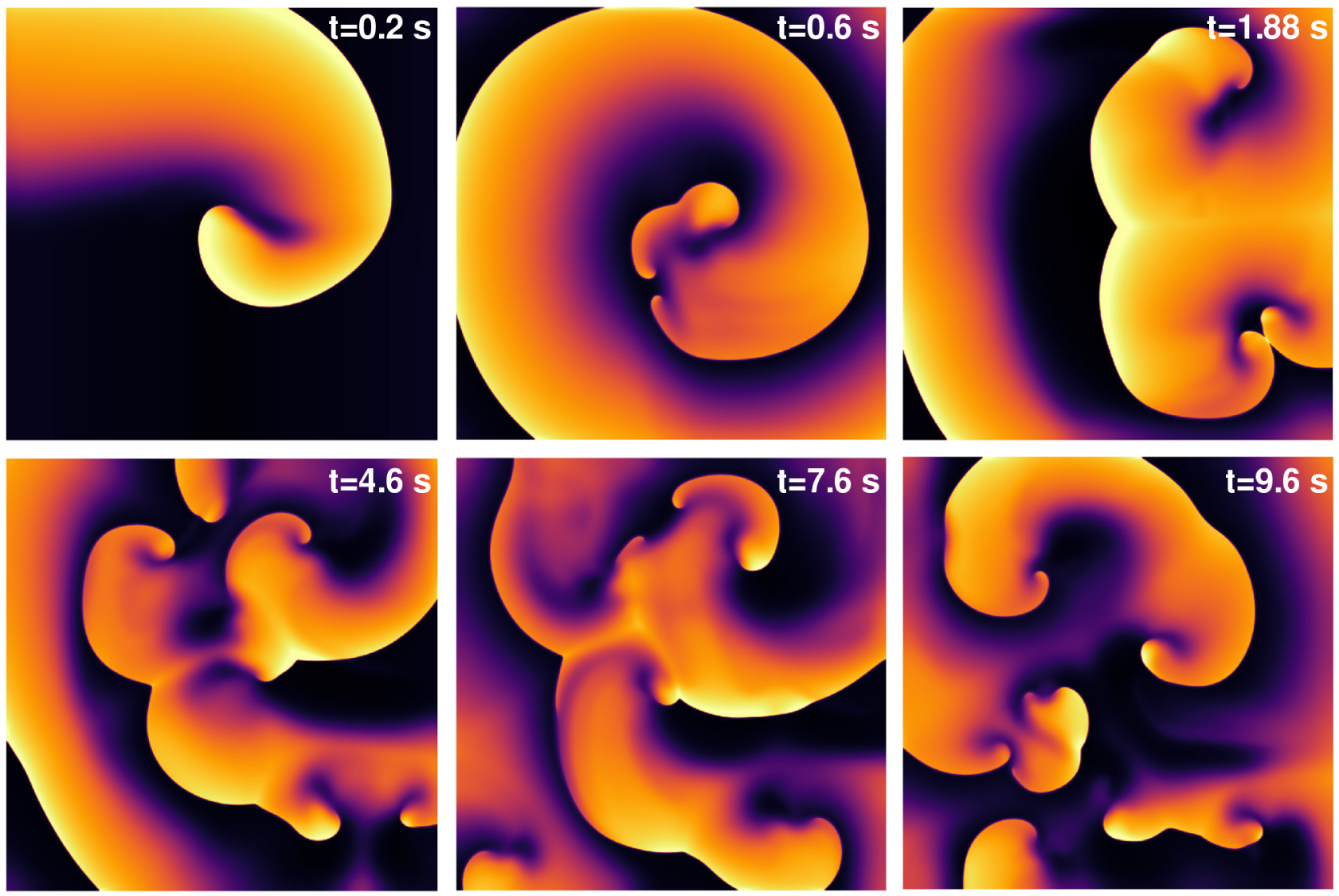
Spiral wave breakup in 2D pig atrial tissue with *G*_*Kr,max*_ reduced to 0.25x its value in the healthy pig model. Pseudocolour plots of the membrane voltage distribution at different times demonstrates the occurrence of multiple spiral waves in the domain.

## Discussion

In this study, we present the first complete mathematical model of the pig atrial tissue. It is built upon experimental data on pig atria as obtained from literature, and new patch-clamp data that was produced in our own laboratory. The model is numerically stable over long timescales, and is capable of reproducing pig atrial action potentials that can be compared closely with experiments. In particular, the AP characteristics, namely APD, CV, RMP, APA and 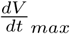 show excellent agreement with experiments, not only for a single evoked AP, but also for the full extent of their respective restitution curves. This confirms that our model is capable of reproducing the exact electrical response as can be expected from healthy pig atrial cardiomyocytes.

Our model takes into consideration the uniqueness of the constitution of the transient outward current. In most mammalian tissue, this current is found to be predominantly *K*^+^ based. However, in pig atria, this current is solely *Cl*^*−*^-based and activated by the flow of *Ca*^2+^ ions. The unique dynamics of this current results in a downward trend of the early repolarisation APD restitution curves; a feature that is not observed in most mammalian species. Our model reproduces this experimental trend in early APD curves for large cycle lengths, and attributes the trend to the inactivation of the *I*_*ClCa*_ at low cycle lengths [17].

In 2D, we demonstrate the ability of the model to sustain stable spiral waves and spiral wave breakup, which adds to the suitability of the model for *in silico* studies of AF in extended media. A Fourier analysis of the tip trajectory shows that there are 4 fundamental frequencies responsible for the dynamics of the intact spiral wave. Of these frequencies, one is associated with wave meander at *f*_*meander*_ ≃ −0.1 *Hz*, two are associated with the hypocycloid pattern, *f*_0_ = 5.426 *Hz* and *f*_1_ = − 3.548 *Hz*, with one of those frequencies also being the frequency of rotation of the spiral arm (and thus setting the average stimulation frequency). Furthermore, our model points to the occurrence of alternans in 2D in the presence of spiral waves, between the cycle lengths of 165ms and 215ms. This may explain the difficulty encountered in experiments, with evoking consistent APs at a pacing frequency of 5Hz. To understand the underlying basis of this alternans, we tested an approach suggested by Gomis-Tena and Saiz [42], who inhibitted the *Ca*^2+^-activated *Cl*^−^ current in their canine model to inhibit alternans. Our model, however, failed to show suppression of alternans by similar inhibition of the *I*_*ClCa*_, suggesting that the alternans was not driven by the *Cl*^−^ current.

The model presented here has the same general limitations as any other ionically-detailed mathematical model of cardiac electrophysiology. As previously discussed by Cherry and Fenton [37], detailed mathematical models need to be treated with extreme considerations to their ability to correctly reproduce phenomena outside the general experimental conditions they were modelled after, and their main utility should be in developing new hypotheses in the study of already-known phenomena, rather than for the study of the dynamics of novel, unverified phenomena.

The model falls prey to the natural variability found in cardiac tissue, especially in the atria. The atrial cavities are particularly complex, more so than the ventricles, when it comes to heterogeneity and anisotropy, and the properties of cardiomyocytes are known to be affected by factors like age or sex [37]. In the context of this project, this is evidently palpable in the description of *I*_*Kur*_ given in the two published papers used as sources in this model, which differ significantly, and it highlights some of the compromises that researchers must make when building a general model.

While the study on spiral tip patterns in this model is thorough, a lack of resolution due to computing limitations means that 1 of the main driving frequencies of the system can only be approximately studied, and another must be ignored altogether. Further simulations with sampling for a much longer period would be needed to solve this resolution problem and map out the 2 smaller frequencies with good enough resolution.

Some of the limitations of the model come from the fact that it relies on experiments from literature for the description of individual currents, which sometimes have incomplete data (lack of information of the time dynamics of the currents in Li *et al*., for example [17]), or which have fundamentally different experimental setups (Li *et al*. vs Ehrlich *et al*. [17, 25]). However, this model provides a good basis to start, and can be develeoped further, as and when new experimental data become available.

One important limitation of the model lies in its description of the Ca^2+^ dynamics, which is mostly taken from the human atrial model of Courtemanche *et al*.. The CRN model itself adapts the description of the *Ca*^2+^ dynamics from the Luo-Rudy model for guinea pig ventricular cardiomyocytes [19]. Thus the *Ca*^2+^-dynamics cannot be called state-of-the-art. Although it does give rise to physiologically relevant pig atrial action potentials, the model does not provide any significant insight to the fundamental role that Ca^2+^ plays in mediating I_ClCa_ (*T*_*to*_) and early AP repolarisation. It would therefore be of great interest to make detailed experimental measurements on Ca^2+^ dynamics specific for the pig atria, with the aim of building a more accurate mathematical description to elucidate the mechanisms underlying the dynamics of *T*_*to*_ and to make more accurate predictions of its behavior in arrhythmias.

## Materials and methods

All animal care and use procedures were carried out exclusively by appropriately trained staff and were in accordance with the German Animal Welfare Act and reported to the local animal welfare officers. The handling of the animals prior to the experiments and the humane, animal welfare procedures strictly followed animal welfare regulations, in accordance with German legislation, local regulations and the recommendations of the Federation of European Laboratory Animal Science Associations (FELASA). All scientists and technicians involved have been accredited by the responsible ethics committee (Lower Saxony State Office for Consumer Protection and Food Safety - LAVES).

## Experimental Recordings

Trabecular muscles were isolated and excised from a whole right atria, and placed in a custom-built recording chamber under continuous perfusion of heated (37°C) and carbonated (5% CO_2_, 95% O_2_) Tyrode’s solution containing (in mM): NaCl 126.7, KCl 5.4, MgCl_2_ 1.1, CaCl_2_ 1.8, NaHPO_4_ 0.42, NaHCO_3_ 22, glucose 5.5, pH=7.45 at least 45 min before measurement, for accommodation.

Borosilicate glass capillaries (Hilgenberg, Germany) were pulled using a horizontal pipette puller (Zeitz, Germany). Electrical resistance was 30-40 MΩ. Pipettes were backfilled with 3M KCl.

Tissues were electrically stimulated with a 1 ms monophasic pulse using a custom-made electrode (FHC, USA). Pulse amplitude was pre-defined as 30% higher than the value necessary to trigger an action potential. After successful tissue impalement, and after reaching steady state activity, the tissue was then subjected to a train of electrical stimulation at increasing frequencies (0.25 Hz, 0.5 Hz, 1 Hz, 2 Hz, 3 Hz and 4 Hz). AP onset at 5Hz proved difficult and inconsistent.

Membrane potential signals were amplified using a Sec-05-X (npi, Germany) amplifier, digitised using LabChart PowerLab, and acquired and saved with LabChart Pro 7 software (both: ADInstruments, New Zealand).

Analysis was performed using LabChart pro and GraphPad Prism 7 (GraphPad Software Inc., USA). The average value of 10 consecutive action potentials were calculated in LabChart Pro. The following parameters were measured: resting membrane potential (RMP), action potential maximum upstroke velocity (dV/dtmax), action potential amplitude (APA) and the action potential duration at 20, 50 and 90% of repolarisation (*APD*_20_, *APD*_50_ and *APD*_90_ respectively).

## Mathematical Model

The fitting of IV curves from experimental data by Li *et al*. [17] was carried out by minimization of the squared error between the simulated and experimental data using Python’s Scipy module [40]. For this purpose, a function was created in Python that would recreate the patch-clamp experiments as in Li *et al*. [17], and output the simulated IV curve. The morphology of each individual current and its gating variables was initially taken directly from the human atrial model by Courtemanche *et al*. [19] and corrected accordingly to match experimental data. Fitting was done by matching normalised IV curves first, and then adjusting conductance values to match the non-normalised experimental IV curves.

Overall AP morphology and restitution curves were later matched by re-adjusting conductance values of the different currents, and simulating AP evolution for stimulation at different cycle lengths. Slight changes were made to currents not fitted from Li *et al*. [17] to better adjust experimental restitution curves. See the Results section for detailed explanations of each individual current.

## Spatial extension of the model to higher dimensions

To simulate wave propagation in 1 dimension and above, we added a diffusion term to Eq. 1, such that the spatiotemporal evolution of the voltage is given by

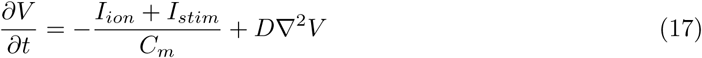

We used a value 0.00126 *cm*^2^*/ms* for the diffusion coefficient *D*. This choice of *D* allowed our model to reproduce the experimentally observed conduction velocity of 58 cm/s from Jang *et al*. [38].

For numerical integration of Eq. 17, we used a Forward Time Centered Space (FTCS) scheme, with a space differential Δ*x* = Δ*y* = 0.022 *cm*. The timestep chosen for the simulations was Δ*t* = 0.02*ms* and all coding was done using Python or C, with MPI-based parallelization.

## Supporting Information

**S1 Appendix Complete set of equations of the pig atrial model**. Includes scheme used for the numerical integration of the model.

**S1 Table Set of model parameters and universal constants. S2 Table Set of initial conditions used in the model**.

**S1 Video**. Pseudocolour plots of the membrane voltage distribution during the formation and spatiotemporal evolution of a spiral wave. The domain contains 512 × 512 grid points and the video plays at 10fps. The total video represents 10*s* of simulation.

**S2 Video**. Pseudocolour plots of the membrane voltage distribution during the breakup and spatiotemporal evolution of a spiral wave in altered parameter regime. Specifically, the maximum conductance for the *I*_*Kr*_ is reduced by a factor of 4. The domain contains 1024 × 1024 grid points and the video plays at 10fps. The total video represents 10*s* of simulation.

## Acknowledgments

VP would like to thank Prof. Blas Echebarria for useful discussions.

## Author Contributions

Designed and developed the model: VP, RM. Conceived and carried out the restitution experiments: TR, FEF, NV. Wrote the paper: VP, RM. Read and reviewed the manuscript: VP, RM, SL, TR, FEF, NV. Acquired funding: SL, NV.

